# Atomic view into *Plasmodium* actin polymerization, ATP hydrolysis, and phosphate release

**DOI:** 10.1101/467423

**Authors:** Esa-Pekka Kumpula, Andrea J. López, Leila Tajedin, Huijong Han, Inari Kursula

## Abstract

*Plasmodium* actins form very short filaments and have a non-canonical link between ATP hydrolysis and polymerization. Long filaments are detrimental to the parasites, but the structural factors constraining *Plasmodium* microfilament lengths are currently unknown. Using high-resolution crystallography, we show that magnesium binding activates the *Plasmodium* actin I monomer before polymerization by a slight flattening, which is reversed upon phosphate release. A coordinated potassium ion resides in the active site during hydrolysis and leaves together with the phosphate, a process governed by the position of the Arg178/Asp180-containing A-loop. Asp180 interacts with either Lys270 or His74, depending on protonation, while Arg178 links the inner and outer domains. Hence, the A-loop is a switch between stable and non-stable filament conformations. Our data provide a comprehensive model for polymerization, phosphate release, and the inherent instability of parasite microfilaments.

---

Actin is the constituent protein of microfilaments with essential roles in central processes in the cell, including transport, cell division, and motility^1–3^. The primary biological activity of actin is its polymerization to form filaments that can generate force at cell membranes or act as scaffolding structures or tracks for motor proteins^4^. These filaments are on a timer, based on the hydrolysis of tightly-bound ATP, formation of the stable intermediate ADP-P_i_-actin, and finally, the release of inorganic phosphate (P_i_)^5^. In model actins, the coupling of nucleotide hydrolysis to filament stability is well established. In general, ADP-actin depolymerizes much faster than ATP or ADP-P_i_ actin and is therefore the main depolymerizing species^6^. Although ADP-actin can polymerize, its critical concentration is much higher than that of ATP-actin^6^, which leads to domination of ATP-actin in polymerization kinetics. Outliers of this functional consensus are actins of the phylum Apicomplexa, including *Plasmodium* spp. and *Toxoplasma gondii* – both notorious human pathogens. Actins of these parasites are among the most evolutionarily divergent eukaryotic actins, while still retaining most of the core features of canonical actins^7–10^. The primary actin of *P. falciparum* and the only one of *T. gondii* are the best understood of the phylum, while others remain virtually uncharacterized.

*In vitro*, apicomplexan actins tend to form only short filaments of ~100 nm without the filament-stabilizing macrolide jasplakinolide^7–9,11^. *T. gondii* actin (TgAct) has been proposed to follow an isodesmic polymerization mechanism^10^, which would differ fundamentally from the classical nucleation-elongation pathway. However, *P. falciparum* actin I (*Pf*ActI) has a critical concentration close to that of mammalian α-actin and a very similar elongation rate^12^. Under ADP-rich conditions, *Pf*ActI forms oligomers of 3-12 subunits, while forming larger polymeric species in polymerizing conditions, together with a significant pool of dimers^8,12^.

Structurally, the *Pf*ActI monomer resembles canonical actins^8^ **(Fig. 1a**). The largest structural differences are at the pointed end, namely subdomain (SD) 2 (containing the DNaseI-binding D-loop) and parts of SD4, which both connect to SD3 of the next longitudinal protomer in the filament. The D-loop and the C terminus are both important functional factors but are disordered in the crystal structure of *Pf*ActI, reflecting their flexibility^8^. In jasplakinolide-stabilized *Pf*ActI filaments, the D-loop is in a clearly altered conformation compared to α-actin filaments^9^. Yet, the main hydrophobic interactions are conserved, and the amino acid substitutions are primarily located at the base of the D-loop^8^. In addition, differences in the plug region (residues Ser266-Ala273, especially Lys270), and some other residues along the filament interface (in particular Val288, Gly200) also likely contribute to filament instability^9^.

**Fig. 1.**
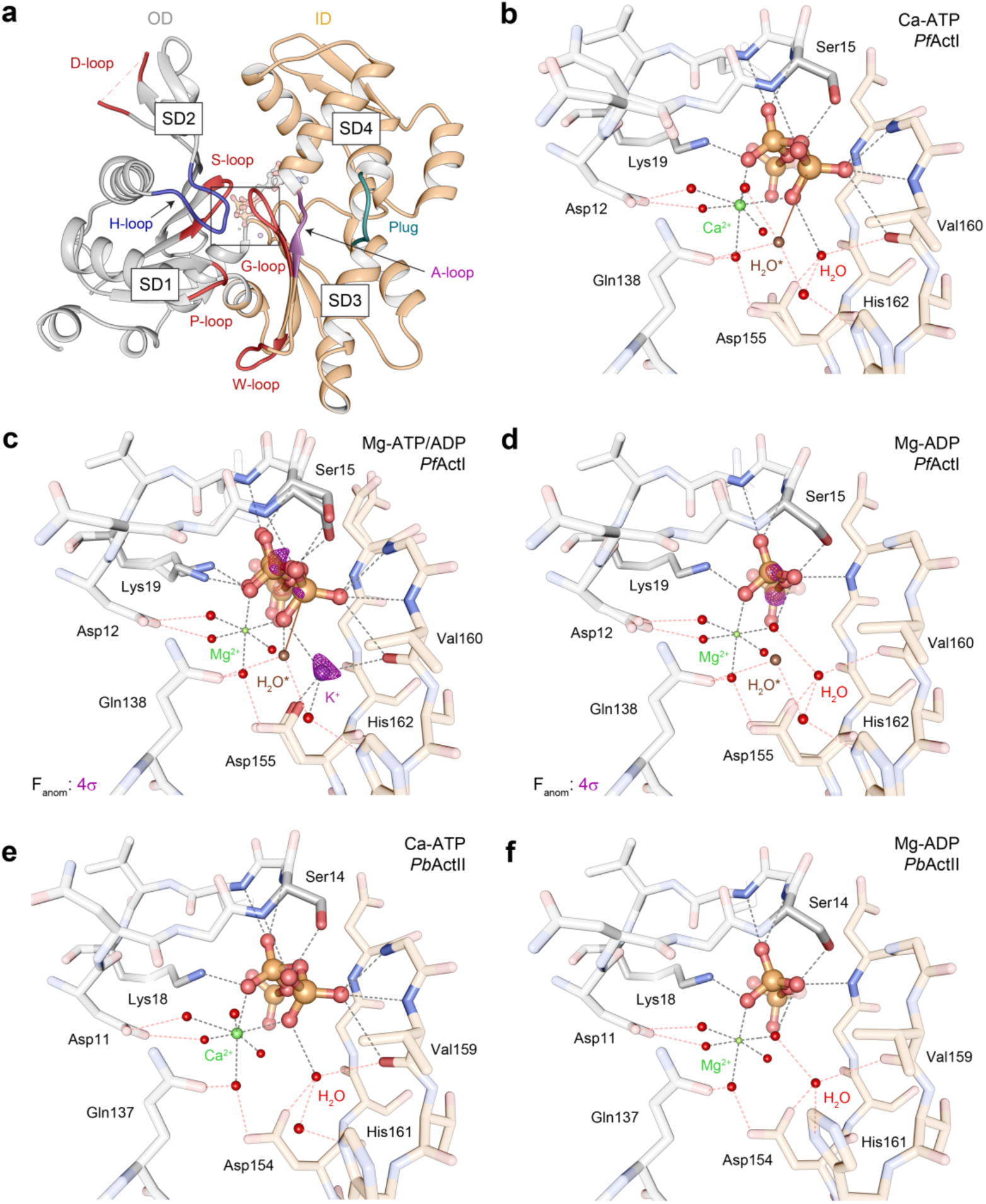
Active site configurations in the *Pf*ActI and *Pb*ActII structures. (a) Overview of the Mg-ATP/ADP-*Pf*ActI monomer with the D-loop, S-loop, H-loop, G-loop, P-loop and W-loop as well as the plug and A-loop indicated. The region of interest enlarged in the other panels is boxed. (b-d) *Pf*ActI structures in the (b) Ca-ATP^8^, (c) Mg-ATP/ADP, and (d) Mg-ADP states. (e-f) *Pb*ActII structures in (e) Ca-ATP^8^ and (f) Mg-ADP states. In all panels, hydrogen bonds with ATP, ADP, or ions are indicated with black dashed lines and the outer shell hydrogen bonding via water molecules with red dashed lines. In (b-c), the brown solid line indicates the nucleophilic attack vector of the putative catalytic water^34^ (H_2_O^∗^). In (c-d), anomalous difference density is shown as a purple mesh at a 4σ contour level. The inner domain (ID) and outer domain (OD) are colored in orange and gray, respectively, in all panels.

The P_i_ release pathway from skeletal muscle α-actin has been studied in detail using molecular dynamics^13^. In the proposed model, the nucleotide is exchanged *via* a so-called front door, where the nucleotide is inserted into the active site, and P_i_ (together with the cation) exits *via* a back door on the opposite side. In a follow-up study, this pathway was determined in detail, and it was suggested that, on its way out, the P_i_ interacts mainly with His73 and Arg177^14^.

Interestingly, it seems that hydrolysis of ATP and subsequent P_i_ release is favorable for oligomerization of *Pf*ActI. Structural changes upon these could thus favor nucleus formation – *i.e*. result in a conformation closer to the filament structure than that of the monomer. Here, we analyze phosphate release rates and high-resolution structures of wild-type and mutant *Plasmodium* actins in different nucleotide states, bridging the gap between structure and function in understanding the polymerization mechanism.

## Results

### Phosphate release is decoupled from polymerization in *Pf*ActI

In skeletal muscle α-actin, conformational changes upon polymerization activate nucleotide hydrolysis in the actin protomers, and the subsequent P_i_ release leads to destabilization of the “aged” filament^15,16^. α-actin releases P_i_ at rates of 0.15-0.47 × 10^-4^ s^-1^ at equilibrium^17^ and 14.8 × 10^-4^ s^-1^ during polymerization^18^ (**Supplementary Table 1**). By comparison, equilibrium P_i_ release rates measured from *Pf*ActI and *Pb*ActII in the Ca state were 1.3 × 10^-4^ s^-1^ for both actins and in the Mg state 3.1 × 10^-4^ s^-1^ for *Pf*ActI and 1.9 × 10^-4^ s^-1^ for *Pb*ActII^8^. These measurements were conducted above the critical concentration of either filament end in the ATP state (1.5 μM for the barbed end, 4.5 μM for the pointed end in 1 mM MgCl_2_^6^). To further characterize the relationship between phosphate release and polymerization, we measured P_i_ release rates from *Pf*ActI, *Pb*ActII and α-actin in 0.2 mM Ca^2+^, 1 mM Mg^2+^, and 5 mM Mg^2+^/50 mM K^+^ at protein concentrations around 1 and 3-6 μM each. Contrary to α-actin, P_i_ release rates of the parasite actins did not increase in the polymerizing MgK conditions at low actin concentrations (**Table 1** and **Supplementary Fig. 1**). This was true also for higher concentrations of *Pf*ActI but not for *Pb*ActII (**Table 2**). At higher concentrations, P_i_ release from *Pf*ActI was instantaneous, while it had a lag phase in *Pb*ActII and α-actin (**Supplementary Fig. 1**). These data suggest that nucleotide hydrolysis and P_i_ release are decoupled from polymerization in *Pf*ActI.

**Table 1:** Phosphate release rates of actins in Ca, Mg and MgK conditions and activation by Mg^2+^ and K^+^ at actin concentrations of 1 μM.

|  | Condition |  |  | Activation |  |
| --- | --- | --- | --- | --- | --- |
|  | Ca<br>(10<sup>-4</sup> s<sup>-1</sup>) | Mg<br>(10<sup>-4</sup> s<sup>-1</sup>) | MgK<br>(10<sup>-4</sup> s<sup>-1</sup>) | by Mg<sup>2+</sup><br>(Mg/Ca) | by K<sup>+</sup><br>(MgK/Mg) |
| K270M<sup>†</sup> | 0.21±0.01<sup>*</sup> | 4.6±0.19<sup>*</sup> | 6.2±0.60<sup>*</sup> | 22±1.7<sup>*</sup> | 1.4±0.2<sup>*</sup> |
| A272W<sup>†</sup> | 0.52±0.02<sup>*</sup> | 9.78±0.06<sup>*</sup> | 9.9±0.19<sup>*</sup> | 18.9±0.9<sup>*</sup> | 1.02±0.03<sup>*</sup> |
| A272C<sup>†</sup> | 0.30±0.01<sup>*</sup> | 1.54±0.06 | 1.69±0.06<sup>*</sup> | 5.1±0.4<sup>*</sup> | 1.09±0.08<sup>*</sup> |
| E49G<sup>†</sup> | 0.58±0.02<sup>*</sup> | 2.8±0.13<sup>*</sup> | 2.8±0.20<sup>*</sup> | 4.7±0.4<sup>*</sup> | 1.0±0.1 |
| F54Y<sup>†</sup> | 1.20±0.08<sup>*</sup> | 3.14±0.09<sup>*</sup> | 2.8±0.12<sup>*</sup> | 2.6±0.3 | 0.90±0.07 |
| P42Q/E49G<sup>†</sup> | 0.52±0.03<sup>*</sup> | 1.18±0.04<sup>*</sup> | 1.18±0.09 | 2.3±0.2 | 1.0±0.1 |
| *PfAct1 wt* | 0.74±0.03 | 1.62±0.05 | 1.3±0.12 | 2.2±0.2 | 0.8±0.1 |
| G115A<sup>†</sup> | 0.70±0.07 | 1.05±0.04<sup>*</sup> | 1.00±0.07 | 1.6±0.6 | 1.0±0.2 |
| P42Q<sup>†</sup> | 1.24±0.03<sup>*</sup> | 1.75±0.02 | 1.52±0.09 | 1.41±0.05<sup>*</sup> | 0.87±0.07 |
| H74Q<sup>†</sup> | 0.27±0.03<sup>*</sup> | 0.27±0.01<sup>*</sup> | 0.21±0.03<sup>*</sup> | 1.0±0.2<sup>*</sup> | 0.8±0.2 |
| *PbActII* | 0.60±0.04<sup>*</sup> | 0.67±0.05<sup>*</sup> | 0.56±0.02<sup>*</sup> | 1.1±0.1<sup>*</sup> | 0.83±0.09 |
| α-actin | 0.22±0.01<sup>*</sup> | 0.68±0.01<sup>*</sup> | 1.99±0.02<sup>*</sup> | 3.1±0.2<sup>*</sup> | 2.92±0.08<sup>*</sup> |

\*: P < 0.01 (N = 3), two-tailed Student's T-test vs. corresponding values of *PfActI* wildtype.
†: Mutants are of *PfActI*.
*PfActI* results are ordered by Mg<sup>2+</sup> activation.
Errors represent standard deviations.

**Table 2.** Phosphate release rates of actins in Ca, Mg and MgK conditions and activation by Mg^2+^ and K^+^ at actin concentrations of 3-6 μM.

|  | Condition |  |  | Activation |  |
| --- | --- | --- | --- | --- | --- |
|  | Ca<br>(10<sup>-4</sup> s<sup>-1</sup>) | Mg<br>(10<sup>-4</sup> s<sup>-1</sup>) | MgK<br>(10<sup>-4</sup> s<sup>-1</sup>) | by Mg<sup>2+</sup><br>(Mg/Ca) | by K<sup>+</sup><br>(MgK/Mg) |
| *PfActI*<br>(3.5 μM) | 0.38±0.004 | 1.7±0.04 | 1.7±0.08 | 4.5±0.2 | 1.0±0.08 |
| *PbActII*<br>(3.8 μM) | 0.15±0.009 | 1.6±0.03 | 7.0±0.4 | 11.0±1.0 | 5.0±0.4 |
| *α-actin*<br>(5.9 μM) | 0.095±0.003 | 2.9±0.06 | 7.0±0.1 | 31.0±1.6 | 2.2±0.09 |

P < 0.01 (N = 3) for all *PbActII* and *α-actin* values, two-tailed Student's T-test vs. corresponding values of *PfActI* wildtype. Errors represent standard deviations.

### Gelsolin-bound *Pf*ActI undergoes slow ATP hydrolysis but fast phosphate release *in crystallo*

Since the major activation of P_i_ release from *Pf*ActI is caused by Mg^2+^, we decided to study the process in detail by analyzing crystal structures of monomeric *Pf*ActI and *Pb*ActII in the Mg state and compare those to the published high-resolution structures of the Ca states^8^. The crystals diffracted to high resolution (1.2-1.85 Å, **Supplementary Tables 2 and 3**), enabling a detailed structural analysis. To our surprise, *Pf*ActI crystals showed a mixture of ATP and ADP in the active site (**Fig. 1**, **Supplementary Fig. 2**, and **Supplementary Text**). Only after aging the crystals for several months, we could obtain data explained by an ADP-only model (**Fig. 1d**). Despite the mixed nucleotide state, we were unable to locate free P_i_ anywhere within the structure, even after soaking the aged crystals in P_i_. Contrary to *Pf*ActI, Mg-*Pb*ActII crystals contained only ADP just two weeks after crystallization, despite showing a slightly lower P_i_ release rate in solution than *Pf*ActI (**Supplementary Fig. 1** and **Table 2**). Thus, the effects of gelsolin and/or the crystalline environment apparently slow down the hydrolysis but not the P_i_ release rate of *Pf*ActI. The combination of high resolution and slow hydrolysis provides a convenient window to visualize the structural changes upon activation of P_i_ release and polymerization.

### ATP hydrolysis in *pf*ActI proceeds through opening and twisting of the monomer

The overall structures of the different nucleotide states of *Pf*ActI appear very similar, but principal component analysis (PCA) with a set of 147 unique actin structures identifies two conformational shifts during the reaction pathway (**Fig. 2, Supplementary Movie 1**): (i) opening of the nucleotide binding cleft and (ii) slight flattening upon inclusion of Mg^2+^, followed by twisting of the monomer upon completion of hydrolysis. A dataset comprising only *Plasmodium* actins shows a similar trend (**Fig. 2c-d**), although PC2 in this dataset depicts a change in SD2 and not so much in SD1, as in the full dataset (**Supplementary Movie 1**). The twist angles of the mass centers of the subdomains (θ) were used as an independent measure and showed angles of 19.0°, 17.9°, and 20.0° for Ca-ATP, Mg-ATP/ADP, and Mg-ADP structures, respectively (**Supplementary Table 4**). The opening-closing motion was not evident from distances of the mass centers of SD2 and SD4 (d_2-4_) or phosphate clamp distances (b_2_) as defined before^19^. However, anisotropic B-factors provided support for the opening, showing a directional destabilization of SD2 towards SD4 (**Supplementary Fig. 3**). A comparable dataset of *Dictyostelium discoideum* actins is characterized in PCA by a combination of opening and twisting upon inclusion of Mg^2+^ and a reversal of the opening upon completion of hydrolysis^20^.

**Fig. 2.**
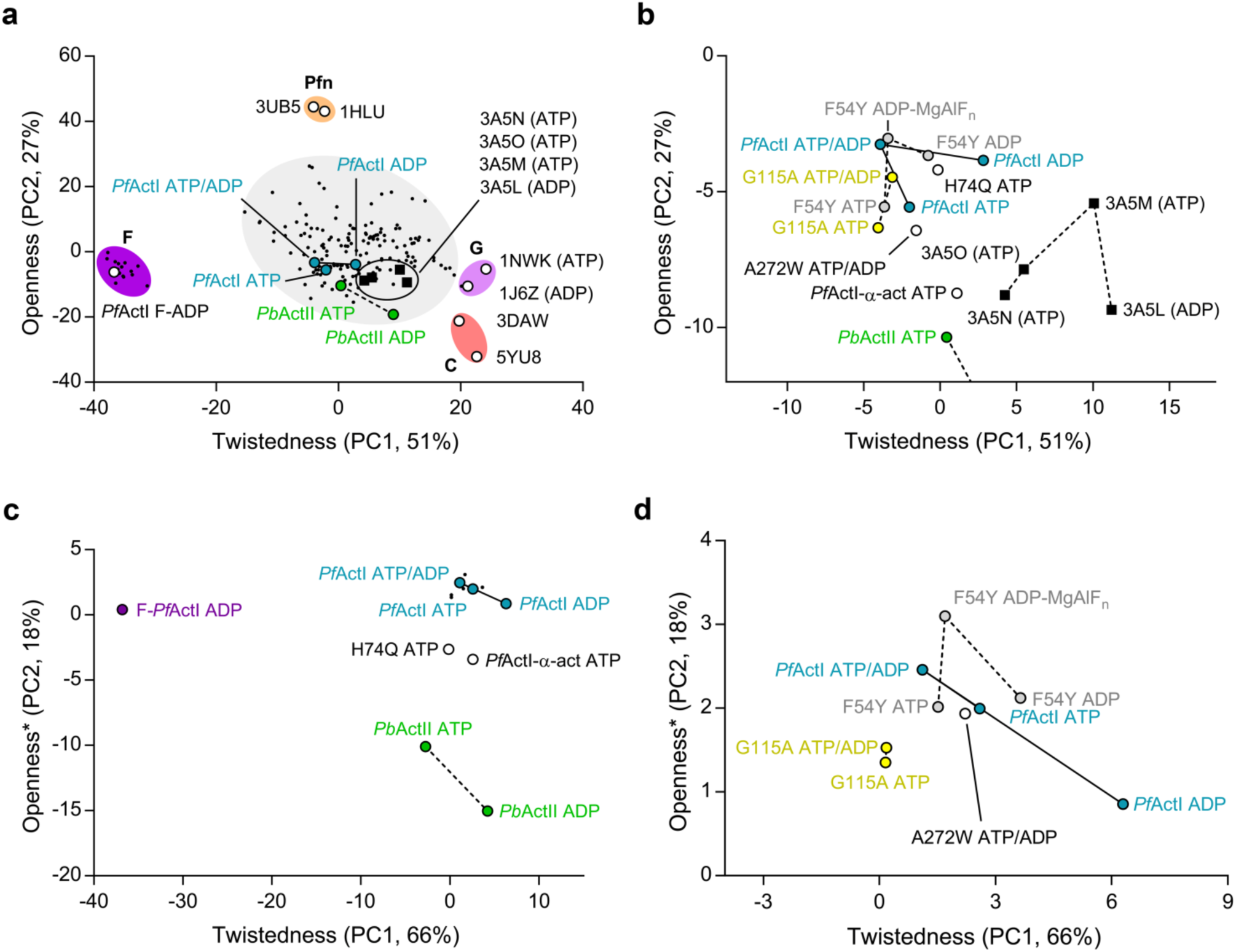
PCA analysis of actins. (a) Plot of twistedness (PC1) vs. openness (PC2) of the full dataset of 147 actin structures (**Online Methods, Supplementary Movie 1**). Defined structural groups of filament structures (dark purple), profilin-bound open structures (orange), free G-actin structures (light purple) and ADF/cofilin bound structures (pink) are indicated with **F**, **Pfn**, **G** and **C**, respectively. The large heterogeneous group in the middle is shaded in gray. Structures of interest are indicated with circles or squares and names or PDB identifiers, whereas others are indicated with black dots. (b) Zoomed view of (a) containing the *Plasmodium* actin structures (excluding *Pb*ActII Mg-ADP) as well as four mutant *D. discoideum* actin structures^20^ constituting a full set of nucleotide and divalent cation states. (c) PCA of *Plasmodium* actin structures only (**Online Methods**), with similar notation as in (a). (d) Zoomed view of (c) containing all relevant *Pf*ActI structures excluding the H74Q mutant and the *Pb*ActI-α-actin chimera^8^. The lines and dashed lines between the *Pf*ActI and *Pb*ActII structures indicate the path in the hydrolytic direction (ATP-ATP/ADP-ADP) as appropriate for each set of structures.

### *Pf*ActI binds potassium during ATP hydrolysis

In the mixed ATP/ADP structure of *Pf*ActI, we identified K^+^ with a final refined occupancy of 0.7, which is close to the occupancy of ATP (0.8), between the side chain of Asp155, the backbone N of Gly157, and the backbone O of Val160 (**Fig. 1c** and **Supplementary Text**). The active site of actin is highly conserved, including the residues coordinating this K^+^. Yet, there is no evidence of K^+^ or any other ions at this site in published actin structures, other than the Cd-ATP-*Pf*ActI structure^21^, where Cd was refined at this site. However, this site corresponds to one of the K^+^-binding sites in the homologous Hsc70 nucleotide-binding domain^22^. The Mg-ADP structure does not contain excess electron density or anomalous difference density at this site (**Fig. 1d**), despite showing anomalous difference density for the Pα and Pβ atoms of ADP. This suggests that K^+^ leaves the active site upon P_i_ release. Since K^+^ does not activate P_i_ release from *Pf*ActI (**Tables 1 and 2**), this interaction most likely does not directly influence the mechanism of P_i_ release in *Pf*ActI but may rather be relevant for hydrolysis.

### Non-methylated His74 and Lys270 play ping-pong on the A-loop in *Pf*ActI

Three loops in the actin fold are considered primary sensors of the nucleotide state (**Fig. 1a**): the S-loop (residues 11-16^23–25^), the H-loop (residues 70-78^24^), and the G-loop (residues 154-161^25^). Other, more distant sensors of the nucleotide state are the W-loop (residues 165-172^26^), the D-loop (residues 38-52^23–25^), and the C terminus (residues 349-375^27^). The foremost nucleotide state sensor in canonical actins is Ser14, whose side chain rotates towards the β-phosphate of ADP upon P_i_ release. The conformation of the corresponding Ser15 in *Pf*ActI moves from the ATP-state^8^ through a double conformation with occupancies 0.8/0.2 in the ATP/ADP-state to a complete ADP-conformation (**Fig. 1b-d**). This conformational switch is further propagated to the flipping of the peptide bond between Glu73 and His74 (**Supplementary Fig. 4**), as seen also in *Pb*ActII and the uncomplexed ATP and ADP structures of several actin structures^20,23,25^.

Asp180 is located in a short loop following β14 (**Supplementary Fig. 5**), sandwiched between the H-loop and the plug residues, including Lys270 (**Figs. 1a and 3**). This loop, which we subsequently call the A-loop, serves as a linker between SD3 and SD4. In the Ca-ATP structure, the A-loop resides close to the H-loop (**Fig. 3a**). Asp180 is in two conformations: either interacting with the Nδ of His74 (3.2 Å, conformation **1a**) or oriented towards Arg178 (conformation **1b**). In the Mg-ATP/ADP structure, the backbone of the A-loop has a second conformation (conformation **2a**) with an occupancy of 0.4 (**Fig. 3c**). In the Mg-ADP structure, only conformations **1b** and **2a** are present at equal occupancies. B-factors match the environment in both Mg structures (**Supplementary Fig. 6**), and the occupancies are in agreement with the estimated protonation state (55%, see **Supplementary Text**) of the histidine side chain. In conformation **2a**, Asp180 forms a salt bridge with Lys270. In conformation **1a**, Asp180 moves to form a salt bridge with His74. Thus, the A-loop is engaged in a ping-pong movement between the two positive charges. Conformation **1b** is analogous to the position of the side chain in the jasplakinolide-stabilized **Pf**ActI filament model (**Fig. 3g**) and in many canonical actin filament models^9,28–30^.

**Fig. 3.**
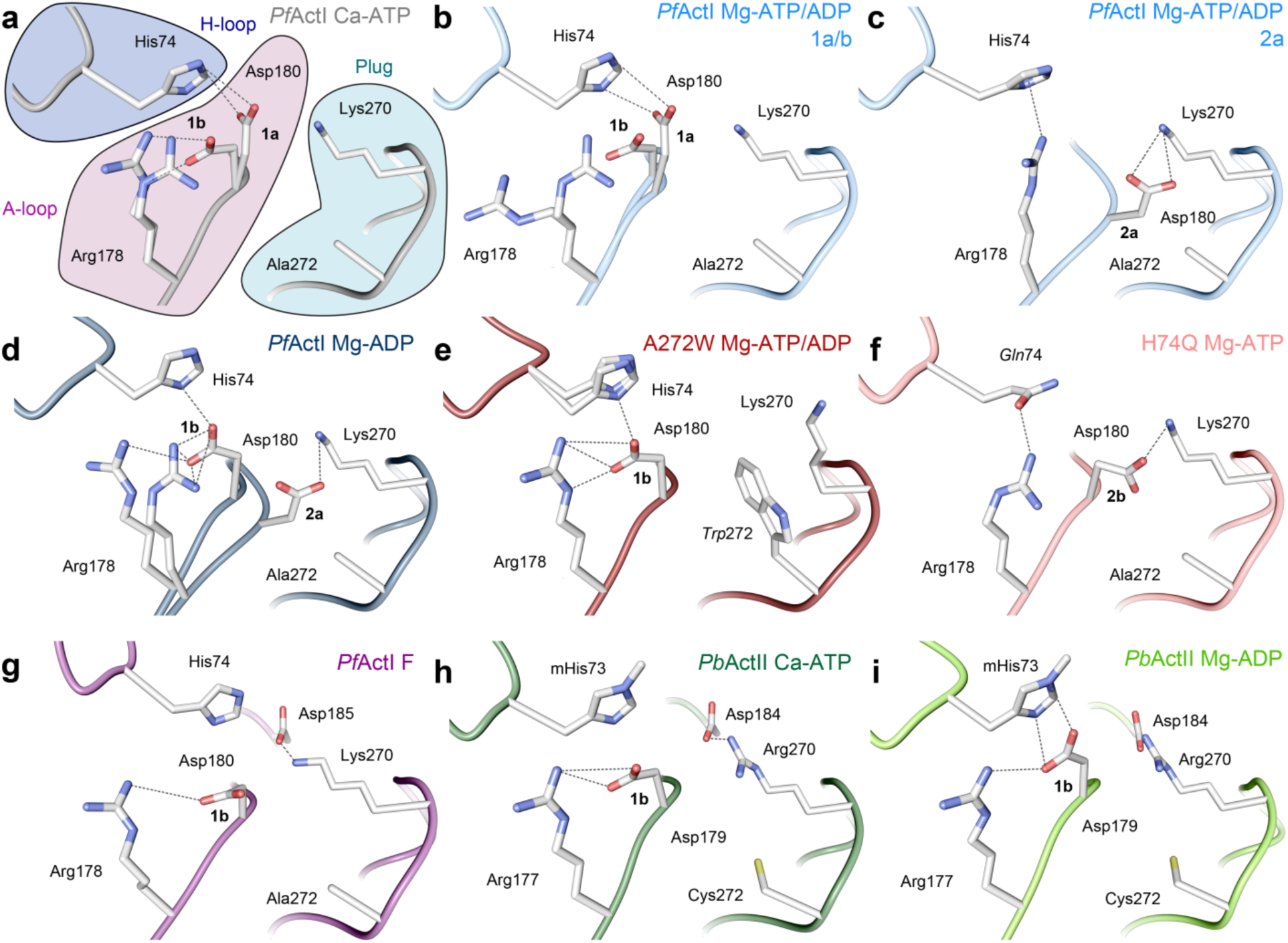
Orientation of the A-loop in *Pf*ActI and *Pb*ActII. (a-d) Wild-type *Pf*ActI in the (a) Ca-ATP state^8^ (**1a** and **1b**), (b) Mg-ATP/ADP state (**1a** and **1b**), (c) Mg-ATP/ADP state (**2a**), and (d) Mg-ADP state (**1b** and **2a**). (e-f) *Pf*ActI mutants (e) A272W in the Mg-ATP/ADP state (**1b**) and (f) H74Q in the Mg-ATP state (**2b**). (g) Wild-type *Pf*ActI in the F-state^9^ (**1b**), stabilized by jasplakinolide (not depicted, see **Fig. 5g** for the jasplakinolide orientation in the filament structure). (h-i) Wild-type *Pb*ActII in the (h) Ca-ATP^8^ (**1b**) and (i) Mg-ADP states (**1b**). In (h), His73 is methylated for consistency even though it is not in the deposited model. The most probable ionic and hydrogen bonding interactions are indicated with dashed lines. The conformers of the H-loop are attributed to each conformation based on overlap of van der Waals radii as well as distances and geometry for hydrogen bonding.

Most model actins, except for that of *Saccharomyces cerevisiae*, presumably have a methylated His74 (*Pf*ActI numbering) in the H-loop^31^, although this is not evident from the majority of structures in the PDB. Our crystal structures are of sufficiently high resolution to verify the previous observations that in native or recombinant **Pf**ActI, His74 is not methylated^11,12^. Curiously, recombinant *Pb*ActII expressed and purified similarly is methylated at this position (**Supplementary Fig. 2**). In actins with a methylated histidine at this site, Nδ is mostly protonated and free to interact with the carbonyl of Gly159 (**Pf**ActI numbering), which together with Val160 is involved in coordinating the active site K^+^ (**Fig. 1c**). As protonated histidines act as cations in electrostatic interactions and as π-systems in cation-π interactions, protonation constitutes a credible interaction switch between His74^+^/Asp180^-^ and His74/Arg178^+^, particularly for a non-methylated histidine (55% protonated at pH 6 based on the typical pK_a_ of histidine side chains in solution). A methylated histidine in canonical actins and *Pb*ActII would favor interactions of the A-loop with the H-loop.

Arg178 in the A-loop participates in connecting the inner (ID) and outer (OD) domains. In the Mg-ATP/ADP structure, Arg178 moves towards the carbonyl groups of His74 and Pro110 in conformation **1b**, thus connecting the P-loop in SD1 (residues 109-114) and H-loop in SD2 (**Supplementary Fig. 7**). Conversely in conformation **2a**, Arg178 interacts with His74 *via* a cation-π interaction, which only maintains the contact between SD3 and SD2. Since the two conformations of the A-loop backbone (**1a/b** and **2a**) are evident in the presence of Mg^2+^ but not with Ca^2+^ and are still present in the Mg-ADP structure, the movement of the loop is either connected directly to Mg^2+^ binding or is an indirect result facilitated by Mg^2+^ binding and the resulting accelerated P_i_ release.

### Structural differences in the Ca and Mg states of *Pb*ActII

According to PCA, Mg-ADP *Pb*ActII is less open and more twisted than the Ca-ATP form, situating towards the twinfilin-C complex^32^ and the cofilin-decorated filament structure^33^. Measurements of θ, d_2-4_ and b_2_ support these findings (**Supplementary Table 3**). However, the largest changes appear in SD2, which has high B-factors and relatively weak electron density (**Supplementary Fig. 9**). The active site configurations in the Ca states are similar between *Pf*ActI and *Pb*ActII (**Fig. 1b, e**). However, in the presence of Mg^2+^, the His161 side chain adopts a different conformation in *Pb*ActII than that seen in any of the structures of *Pf*ActI and most other gelsolin-bound structure in the PDB (**Fig. 1f**). The exception to this is the *D. discoideum* actin structure in the presence of Li-ATP (1NMD), where a similar conformation was proposed to be more amenable to hydrolytic activity^34^. However, the side chain is rotated 180° about the Cβ-Cγ bond in 1NMD compared to *Pb*ActII and most other actin structures. The new conformation of His161 in *Pb*ActII changes the water network by occupying the space of one of the waters coordinating the active site K^+^ in *Pf*ActI. In F-actin, His161 adopts a conformation similar to that seen in *Pb*ActII but even closer to Pγ^30,35^.

There is no evidence of conformations **1a** or **2a** in the *Pb*ActII Mg-ADP structure (**Fig. 3h-i**). This can be rationalized as follows: (i) methylation of His73 ensures that it is mostly protonated and therefore repels Arg177, (ii) Gly115 of *Pf*ActI is threonine in *Pb*ActII, and the G115A mutant also lacks conformation **2a** (see below), and (iii) Ala272 of *Pf*ActI is cysteine in *Pb*ActII, which may sterically block the backbone position of conformation **2a**. The fact that the alternative conformations of the A-loop have not been built in the majority of actin structures does not unambiguously prove that they would not exist, and indeed in several cases, this loop has high B-factors. However, based on available data, we expect that a stable conformation **2a** may be unique to *Pf*ActI, and that *Pb*ActII resembles canonical actins in this respect.

### Canonical-type K270M mutation in *Pf*ActI hyperactivates phosphate release and stabilizes filaments

We proposed earlier that differences in the plug region and especially Lys270 (corresponding to Met269 in α-actin) are among the determining factors for *Pf*ActI filament instability^9^. As Asp180 interacts with Lys270 directly, we generated a canonical-type K270M mutant. Indeed, this mutant formed many more long filaments in the absence of jasplakinolide than wild-type *Pf*ActI (**Supplementary Fig. 9**). Curiously, considering this stabilizing effect, the K270M mutation caused hyperactivation of the P_i_ release rate by Mg^2+^. This activation effect was manifested by a reduction of the rate in Ca conditions to α-actin levels and a moderate increase in Mg. Furthermore, in contrast to the wild type, K270M was no longer insensitive to K^+^ (**Table 1**) and also showed a lag phase at high concentration (**Supplementary Fig. 10a**), thus behaving essentially as α-actin but with a faster rate in Mg and MgK conditions. As this mutation should make conformation **2a** less favorable by disrupting the interaction with Asp180, these results can be taken as indication that conformation **2a** is counterproductive to P_i_ release.

### Mutations affecting the conformational space of the A-loop affect phosphate release in *Pf*ActI

As the A-loop moves into conformation **2a** to interact with Lys270, it fills a space otherwise occupied by water molecules. On the opposite side, Ala272 points towards the A-loop (**Fig. 3a-g**). This alanine is conserved in TgAct and in nearly all alveolates, but is replaced by serine in most model actins and by cysteine in *Pb*ActII or asparagine in *Arabidopsis thaliana* ACT1 (**Supplementary Fig. 5**). We reasoned that if the disappearance of the positive charge by the K270M mutation changed the P_i_ release dramatically, P_i_ release might be directly related to the conformation of the A-loop. Thus, large side chains at position 272 that affect the movement of the A-loop should also modulate the P_i_ release rate. We therefore prepared A272C and A272W mutants - the first to provide a side chain of moderate size, also mimicking *Pb*ActII, and the second to block the movement of the loop completely, both presumably favoring conformation **1a/b**. The A272C mutant caused a moderate 5.1-fold activation upon Mg^2+^ binding, while the A272W mutant showed a large 18.9-fold activation and the largest observed rate (9.78±0.06 × 10^-4^s^-1^) in Mg conditions (**Table 1**).

The A272W structure in MgK conditions resembles overall the mixed structure (RMSD(Cα) = 0.269) more than the Mg-ADP structure (RMSD(Cα) = 0.410) and is positioned close to the Ca-ATP structure in the PCA analysis. The A-loop is forced into conformation **1b** by the Trp272 side chain (**Fig. 3e**). Glu73 is in a double conformation, one similar to the Mg-ADP structure and another to that of the Mg-ATP/ADP structure (**Fig. 3e, Supplementary Fig. 4**). In addition to limiting the conformational space of the A-loop, Trp272 forces Lys270 away from the Asp180 side chain and towards the solvent, widening the gap between His74 and Lys270 from 7.7 to 10.4 Å and only slightly altering the conformation of residues Leu268-Asn281 (RMSD = 0.27 Å, Mg-ATP/ADP-*Pf*ActI compared to Mg-ATP/ADP-*Pf*ActI-A272W) (**Fig. 3e**). The occupancy of ATP in the active site of this relatively fresh crystal is only 0.3 (**Supplementary Table 2**).

To generate a mutant that would favor conformation **2a** of the A-loop, we further prepared a neutralizing H74Q mutant, which negates the charge on the histidine side chain, forcing an unfavorable interaction of the glutamine with Asp180. This mutant was severely compromised in terms of P_i_ release, with α-actin levels of P_i_ release in the Ca state (0.27±0.03 × 10^-4^s^-1^) and no activation by either Mg^2+^ or K^+^ or by using a higher protein concentration (**Table 1**). In this mutant (MgK conditions), the Asp180 side chain is oriented away from Gln74, which interacts with Arg178 (**Fig. 3f**). However, the backbone of the loop did not adopt conformation **2a**, and we therefore call this conformation **2b**, since the carboxylic acid group of the Asp180 side chain occupies the same space as that in conformation **2a**, preserving the interaction with Lys270 (**Fig. 3f**).

### Arg184 interactions with the H-loop in subdomain 2

Interactions across the interdomain cleft mediate twist angle stability and the openness of actin^36^. Upon ATP hydrolysis in *Pf*ActI, Glu73 in the H-loop undergoes a conformational shift, whereby the backbone is flipped and the sidechain orients towards the ID and interacts with Arg184 (**Fig. 4b-d**, **Supplementary Fig. 4**). This conformational shift happens also in *Pb*ActII (**Fig. 4h-i**) and in several canonical actin structures^20,23,25^. In Ca-ATP-*Pf*ActI, Arg184 is engaged in a cation-π interaction with Tyr70. This interaction is preserved in the mixed structure, but is dissipated in the pure Mg-ADP state (**Fig. 4b-d**), after an interaction transfer of Arg184 from Tyr70 to the flipped backbone carbonyl of Glu73. In F-*Pf*ActI, the interaction between Arg184 and Glu73 is enhanced by a hydrogen bond between Arg184 and the Ile72 carbonyl. In the *Pf*ActI H74Q and A272W mutants, the conformations in this area resemble those of the Ca-ATP (in H74Q) and Mg-ADP (in A272W) states (**Fig. 4f-g**, **Supplementary Fig. 4**).

**Fig. 4.**
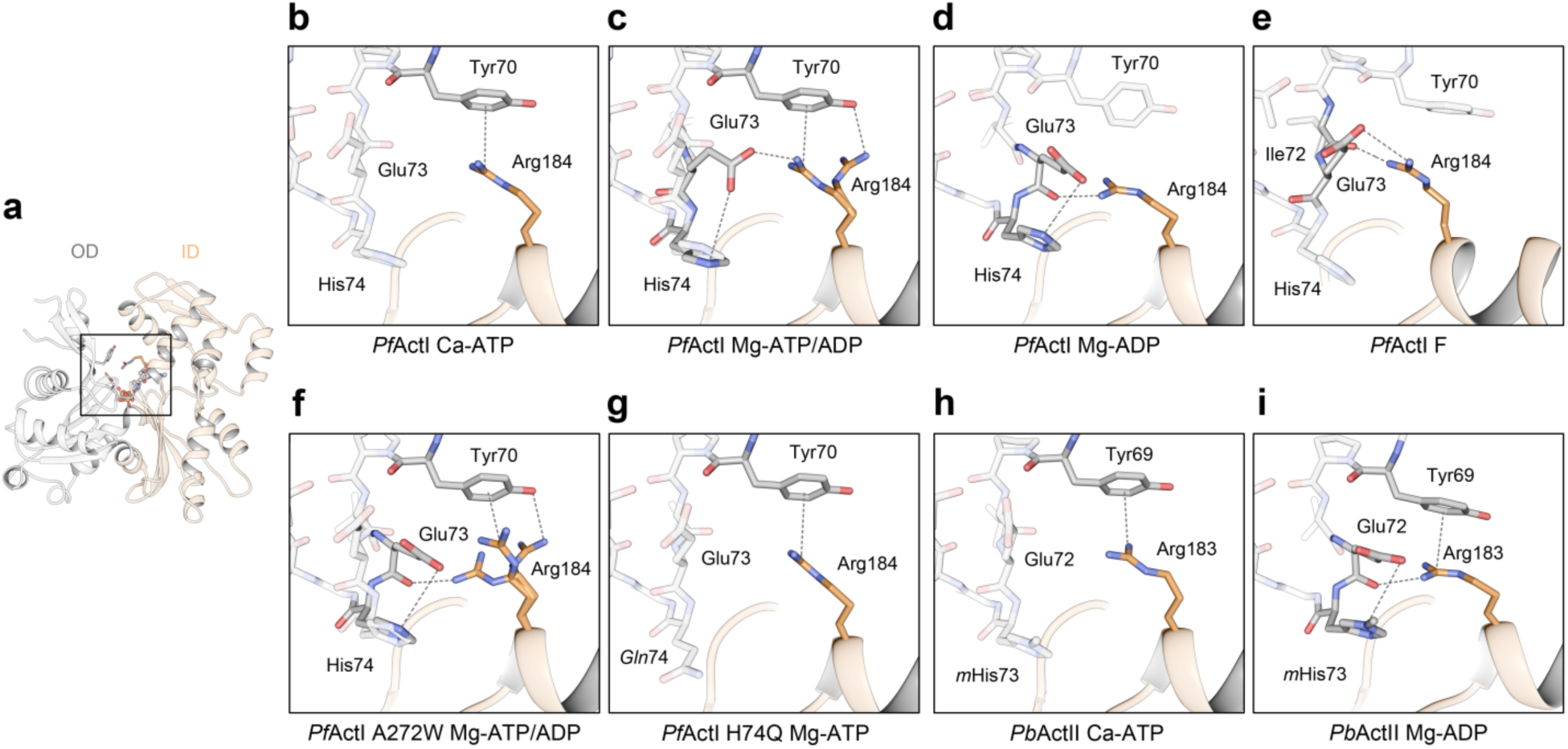
Conformation of the H-loop residues 70-74 as well as the domain cleft spanning Arg184 in *Pf*ActI and corresponding residues 69-73 and Arg183 in *Pb*ActII. (a) Overview of the wild-type *Pf*ActI monomer in the Ca-ATP state^8^ for positional reference. (b-e) wild-type *Pf*ActI in the (b) Ca-ATP state^8^, (c) Mg-ATP/ADP state, (d) Mg-ADP state, and (e) F-state^9^. (f-g) *Pf*ActI mutants (f) A272W in the Mg-ATP/ADP and (g) H74Q in Mg-ATP states. (h-i) Wild-type *Pb*ActII in the (h) Ca-ATP^8^ and (i) Mg-ADP states. The inner domain (ID) and outer domain (OD) are colored in orange and gray, respectively, in all panels. His73 of *Pb*ActII in (h) is methylated for consistency even though it is non-methylated in the original PDB entry. Interatomic distances amenable to ionic interactions or hydrogen bonding are shown as dashed lines.

### The effects of canonical-type mutations in the D-loop on phosphate release

The major substitutions in the D-loop of *Pf*ActI are Pro42, Glu49, and Phe54 (Gln41, Gly48, and Tyr53 in α-actin) (**Supplementary Fig. 5**). Tyr53 is a conserved phosphoregulation site in canonical actins^37^, while the other two sites are interesting because of their possible conformational effects. These residues are invisible or only barely visible (in the case of Phe54) in the crystal structures. However, in the filament, the tip of the D-loop of *Pf*ActI differs from canonical actins^9^. We therefore measured P_i_ release rates for the mutants F54Y^8^, P42Q, E49G, and the double mutant P42Q/E49G of *Pf*ActI. P42Q and E49G showed opposite effects in Mg^2+^ activation with P42Q reducing and E49G increasing it, but both were similarly insensitive to K^+^ (**Table 1**). However, the negative effect of P42Q is due to an increase in the Ca rate compared to wild type, while the positive effect of E49G on Mg^2+^ activation is caused by both reduced rate in Ca and an increased rate in Mg. The double mutant has reduced Mg^2+^ activation with levels indistinguishable from wild-type, while still remaining insensitive to K^+^. Thus, it seems to be dominated by the effect of E49G in the Ca state and shows a compounded negative effect that is not shown by either of the mutations alone.

At high concentration (10 μM), the F54Y mutation reduces the rate of hydrolysis in the Ca state to α-actin levels^8^. Here, we measured the rates at a concentration of 1 μM. The Mg^2+^ and K^+^ - activation levels of F54Y were similar to wild type, but the absolute rates were approximately doubled (**Table 1**). In the Ca condition, the F54Y mutant behaves similarly to P42Q (**Table 2**), while the rates in the Mg and MgK conditions were most similar to the E49G mutant. Thus, these canonical-type mutations in the D-loop area all have similar effects on P_i_ release. However, whereas P42Q and E49G directly affect the tip of the D-loop in the filament, F54Y presents no foreseeable structural changes besides the added H-bonds to Lys62 of monomer n and to Tyr170 of monomer n-2.

### G115A mutation structures the C terminus of *pf*ActI

Gly115 in *Pf*ActI is located in the P-loop of SD1 and is Thr/Ser/Ala in other reference actins (**Supplementary Fig. 5**). Nearby, Pro110 interacts with Arg178 in conformation **1b** and the backbone flexibility conveyed by Gly115 could control the positioning of this interaction. We previously generated a mutant G115A that did not rescue long filament formation in the absence of jasplakinolide but showed slightly longer filaments than wild type in its presence^8^. We crystallized the mutant using the same conditions as the wild-type *Pf*ActI with either Ca^2+^ or Mg^2+^ to compare these structures. Unlike the wild-type (**Fig. 5a**), the C terminus of G115A is more ordered, with interpretable electron density up to Cys375 in the Ca^2+^ and up to His372 in the Mg^2+^ structure (**Fig. 5b-c**). In contrast, all other structures of *Pf*ActI with the exception of H74Q and the *Pb*ActI-α-actin D-loop chimera^8^ have a disordered C terminus after Ser366.

**Fig. 5.**
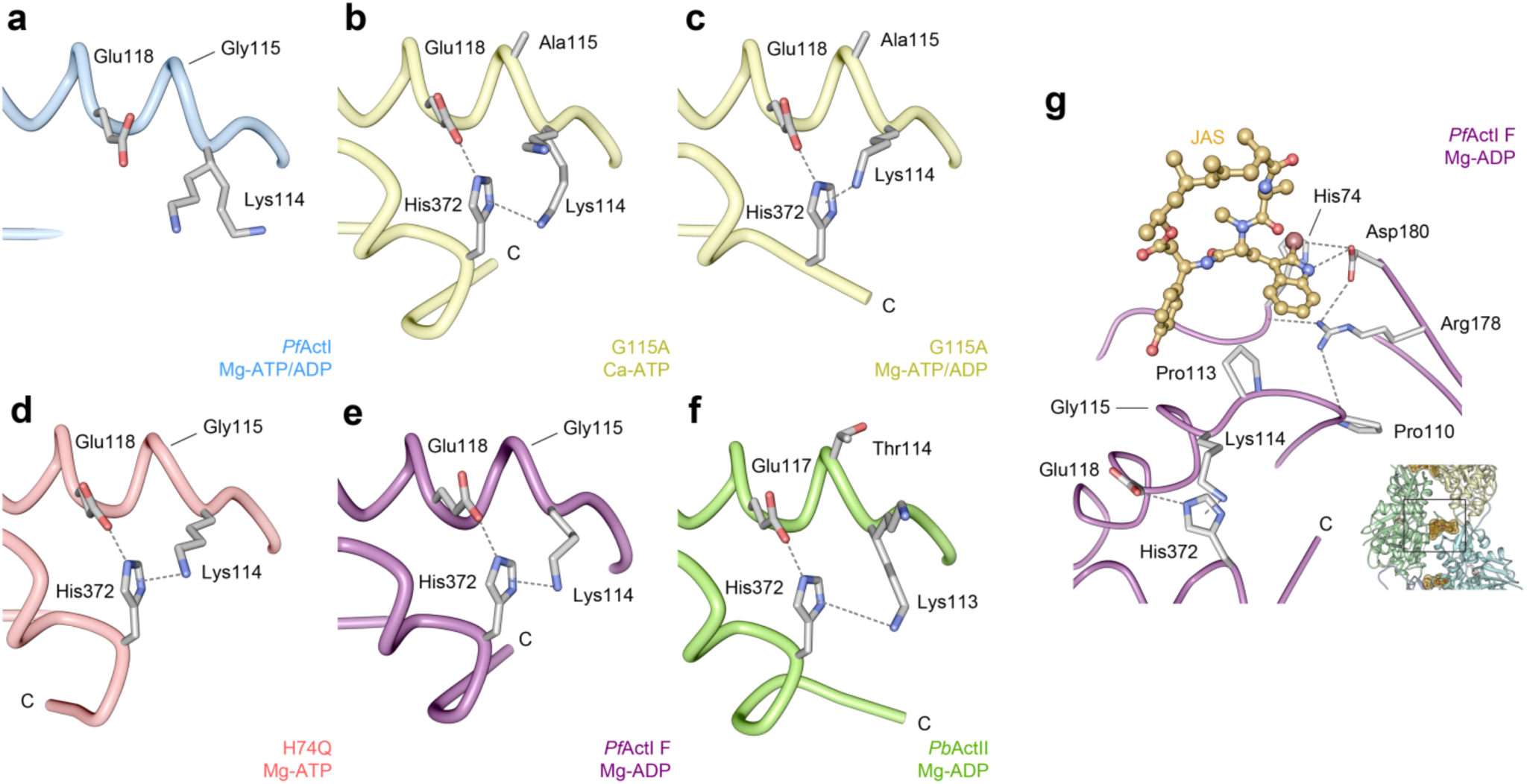
Interaction of the C terminus of *Pf*ActI and *Pb*ActII with Lys114 (Lys113 in *Pb*ActII) and Glu118 (Glu117 in *Pb*ActII) of α3. (a) Wild-type *Pf*ActI in Mg-ATP/ADP state shows a disordered C terminus. *Pf*ActI G115A mutant, in contrast, shows an ordered C terminus in (b) Ca-ATP state and (c) Mg-ATP/ADP state, similar to (d) H74Q mutant in the Mg-ATP state. (e) Wild-type *Pf*ActI in the jasplakinolide-stabilized F-state state^9^ and (f) *Pb*ActII in the Mg-ADP state also have stabilized C termini. The C-terminal His372 interacts with Lys114 and Glu118 of α3 due to the displacement of the N-terminal tip of the helix. In G115A, this is caused by the altered backbone conformation. In H74Q, the effect is likely indirect. In wild-type *Pf*ActI, the C terminus is not stabilized in any gelsolin-bound structure by the His372 interactions, but they are retained in the jasplakinolide-stabilized filament structure due to interactions of the P-loop with the bromoindole moiety in jasplakinolide. In *Pb*ActII (f), residue 114 (corresponding to Gly115 in *Pf*ActI) is threonine and elicits a stabilization of the C terminus. (g) Jasplakinolide interactions with the Proline-rich loop and the A-loop in the filamentous *Pf*ActI structure^9^. Interaction distances amenable to ionic or hydrogen bonding interactions (≤ 4 Å) are indicated with dashed lines. JAS: jasplakinolide, C: C-terminus. The inset in (g) shows the position in the filament.

The G115A mutation straightens α3 and moves the P-loop slightly away from the C terminus. This in turn favors a cation-π interaction between Lys114 and His372 (3.7 Å) and a hydrogen bond between Glu118 and His372 (2.8 Å). In wild type, the position of Lys114 does not allow both interactions to take place simultaneously, which is the likely reason for the disordered C terminus (**Fig. 5a**). In filaments, this interaction is preserved with corresponding distances of 3.1 Å (Lys114-His372) and 3.0 Å (Glu118-His372) (**Fig. 5e**). *Pb*ActII, which has an ordered C terminus in both Ca and Mg states (**Fig. 5f**) has a threonine in the corresponding position 114. The distances from Lys113 and Glu117 to His372 are 2.7 Å and 5.0 Å in Mg-ADP-*Pb*ActII. The altered position of the P-loop does not extend to Pro110, and therefore does not directly influence the interactions of Arg178 at the interface of SD1 and SD2. Trp357 and Phe353 are in a double conformation in both structures, the former facilitating a recently identified cation binding site^38^. The conformations **1a** and **1b** of the A-loop are evident in these structures, but conformation **2a** is not visible in the Mg^2+^ structure. G115A has only slightly decreased P_i_ release rates in Mg and MgK conditions (**Table 1**).

## Discussion

The large-scale conformational transition of the actin monomer from globular to filamentous form has been described from a series of high-resolution filament structures^15,16,20,39^. However, experimental evidence on what exactly triggers the structural transition and the subsequent activation of hydrolysis is still lacking. Key questions are: (i) Why does Mg-ATP actin polymerize more readily than Ca-ATP actin or Mg-ADP actin? (ii) What is the role of K^+^ in polymerization and ATP hydrolysis? Unlike the most studied model actins, *Pf*ActI forms short oligomers also in classical non-polymerizing conditions in the presence of ADP and, on the other hand, stable dimers, in addition to short filamentous structures, in polymerizing conditions^8,12^. The filaments can be stabilized by jasplakinolide, such that a near-atomic resolution structure of the *Pf*ActI filament has been solved, providing hints to the structural features responsible for the inherent instability of the filaments^9^.

All structures reported here and our earlier *Pf*ActI Ca-ATP structure were obtained with K^+^ in the crystallization solution, which provided direct evidence of Mg^2+^ -dependent K^+^ binding in the active site of *Pf*ActI. This is, to the best of our knowledge, the first experimental evidence of K^+^ in the active site of actin. The presence of K^+^ is in conjunction with the Mg-ATP state but not with Ca-ATP or Mg-ADP states. Thus, K^+^ seems to be involved in hydrolysis and leave the active site together with the P_i_. Mg^2+^ binding in the presence of K^+^ causes a slight flattening and opening of the *Pf*ActI monomer, followed by a closing and twisting back upon hydrolysis. This slightly flattened conformation could be the explanation why Mg-K-ATP actin is the fastest polymerizing actin species^6^. Conversely, Mg-K-ADP actin polymerizes weakly in canonical systems^6^, and the twisting upon ATP hydrolysis, as seen for *Pf*ActI, may explain this. However, since the path of the G-F transition may have major intermediates that are off the linear path and cannot be captured by crystallographic analysis, the validity of the connection between polymerization propensity and twist of a G-actin structure remains to be confirmed. It should also be noted that the response of *Pf*ActI to ADP differs from canonical actins^8^.

A structural homolog of actin, Hsc70, has a conserved K^+^ binding site at the same location as *Pf*ActI^22^. The activity of Hsc70 decreases slightly in the presence of ammonium^40^, which is in line with our previous finding that CH_3_COONH_4_ is able to “protect” *Pf*ActI from oligomerization, which in turn is dependent on ATP hydrolysis^8,12^. However, since *Pf*ActI did not respond to K^+^ in P_i_ release assays, the exact role of the active site K^+^ in P_i_ release remains to be investigated. The positive charge on the K^+^ may play a role in orientation of the γ-phosphate or the catalytic water or charge complementation of its conjugate base OH^-^ in the reaction pathway, as has been suggested for Hsc70^41^. Unlike Hsc70 however, the presence of K^+^ is not mandatory for hydrolysis in *Pf*ActI. Yet, its presence may challenge previous hydrolysis mechanisms proposed based on simulations^42,43^.

The interplay between the H-loop, the A-loop and the plug is complex, but our data provide important insights into how the movement of this triad connects to the mechanism of P_i_ release and polymerization. P_i_ release is strongly influenced by the conformational distribution of the A-loop into the two configurations **1a/b** and **2a/b** as we show by P_i_ release measurements (**Tables 1 and 2**) and structures (**Fig. 2**). Conformation **2b** is counterproductive to P_i_ release, while elimination of **2a** by steric hindrance (as in the mutants A272W and A272C) or by charge neutralization (K270M) favors P_i_ release, suggesting that interactions of the A-loop with the H-loop and the P-loop are required for native activity levels. *In vivo*, mutation K270M is lethal in the blood stages of parasite life cycle^44^. Methylation of His73/74 and the resulting change in side chain charge distribution is a key modulator of P_i_ release. A methylated histidine, as found in most actins, is ~11-fold less protonated in the cellular pH than a non-methylated histidine would be. The only other species with a non-methylated histidine at this position and for which there are structures available is *S. cerevisiae*, which like *Pf*ActI has a shorter lag phase of polymerization and no lag in phosphate release upon polymerization^45^. However, in ScAct, conformation **2a/b** is not present, possibly due to the presence of Leu269 and Ala114 instead of Lys270 and Gly115^46^.

F-actin-like interactions in the activated Mg-ATP state can be considered favorable for polymerization. We consider interactions spanning the cleft between ID and OD on the back face of the monomer the most favorable for flattening and therefore nucleation and polymerization. There are only two such interaction sites: (i) between Arg184 of SD4 and Tyr70 and Glu73 of SD2 (**Fig. 3**) and (ii) between Arg178 of SD3 and Pro110 and His74 of SD1 and SD2, respectively (**Supplementary Fig. 8**). In (i), the interaction of Arg184 via a cation-π interaction to Tyr70 is supplemented by an ionic bond with Glu73 in the Mg-ATP/ADP structure, followed by pulling of the Glu73 towards SD2 and a consequent hydrogen bond to the backbone of Ile72 in the F state. Yet, while the polymerization rate of the β-actin R183W mutant was significantly decreased^36^, the α-actin R183G mutant displayed unaltered polymerization kinetics^47^. In (ii), the Arg178 interaction is absent in Ca-ATP actin, but present in conformation **1b** of Mg-ATP/ADP-*Pf*ActI. The interaction is preserved between His74 and Arg178, and further strengthened by hydrogen bonding to the carbonyl of Leu111. An R177H mutant in yeast actin results in an extended lag phase in polymerization^48^, which corroborates that this interaction promotes nucleation. Arg177 is also the site for polymerization-inhibiting ADP-ribosylation by iota toxins^49,50^.

The two substitutions in D-loop (Pro42 and Glu49 in *Pf*ActI) provide one explanation to the unstable nature of *Pf*ActI. These mutations favor the unstable closed D-loop conformation^30^ to such an extent that even in the presence of jasplakinolide, which forces the stable open D-loop conformation in α-actin, the *Pf*ActI filament adopts the closed conformation^9^. Pro42 and Glu49 are in close proximity to the stiffness and polymerization cation sites^51^, which in turn are close to two substitutions in *Pf*ActI, namely Gly200 and Phe54. Together they participate in a complex interplay that is likely one of the major components of filament instability in *Pf*ActI. As P_i_ release of E49G is activated by Mg^2+^ 2.2-fold more than wild type, while the activation of P42Q/E49G and P42Q is equal or less, respectively, one can conclude that these mutations are complementary to each other and that conformational restrictions of the D-loop and P_i_ release rates are reciprocally connected. Like K270M, the mutation P42Q is lethal *in* vivo^44^. Additionally, the effects of the F54Y mutation on overall rates (but not the activation) show that this mutation has a role beyond post-translational modifications. Interestingly, structural information on P_i_ release seems to be “erased” from α-actin filaments by jasplakinolide, which is attributed to the D-loop conformation^30^. The success in preparing *Pf*ActI filaments for cryo-EM by adding jasplakinolide into filaments after polymerizing to equilibrium^9^ shows that the direct binding effects of jasplakinolide can overcome the effects of the constantly closed conformation of the D-loop.

Apicomplexan microfilaments are short but display a relatively normal critical concentration of polymerization, which means that the filament length distribution must result from the overabundance of nucleation, fragmentation, or both. Since the lag phase is very short^12^, increased nucleation likely contributes. However, we believe fragmentation is also important. The K270M mutant forms long helical filaments, releases phosphate quantitatively faster and qualitatively similar to α-actin, and importantly, disfavors conformation **2a** of the A-loop. This conformation is not seen in the filament model, likely because jasplakinolide binds both Arg178 and Asp180, fixing them in a stable conformation. In its absence, the filament structure would permit this conformation. Based on our observations, we propose a model for *Pf*ActI fragmentation (**Fig. 6**). In this model, conformation **2a** in the naked *Pf*ActI filament severs the contact between the ID and OD, leading to destabilization of the monomer twist and filament contacts, eventually causing a break in the filament. The model provides an alternative, perhaps complementary explanation to the electrostatic effects we presented based on the filament model^9^ and would explain the increased pelleting of native *Pf*ActI at low pH^7^. There are no known actin-binding proteins that can directly affect this region of the filament, suggesting that this mechanism could be a major intrinsic determinant of filament lengths *in vivo*. Importantly, while less favorable due to increased protonation of the methylated His73, the lack of attraction between Asp179 and Met269 and the apparent absence of conformation **2a** caused by the G115A substitution, the proposed mechanism could work also in canonical actins. As crystal structures represent low-energy states, it is possible that fragmentation in canonical actins proceeds through the same mechanism, simply less frequently.

**Fig. 6.**
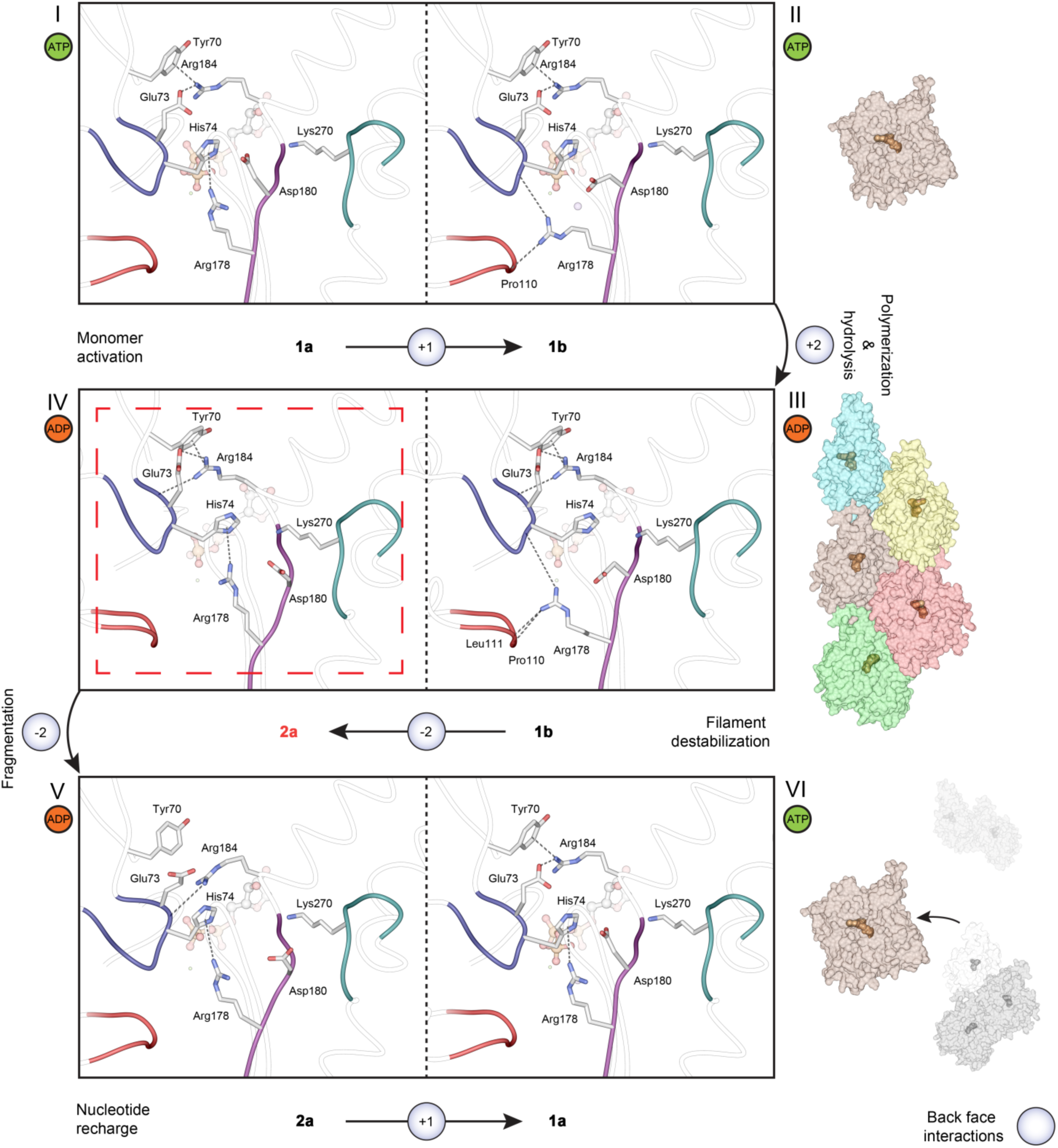
Mechanistic model of **Pf**Actl monomer activation, fragmentation, and nucleotide recharge. Monomer activation takes place by a conformational change from **1a** (I) to **1b** (II), conferring two new back-face interactions that stabilize an F-like conformation. Upon polymerization (III), two new interactions are formed, further stabilizing the flat conformation. In F-*Pf*ActI, ATP is hydrolyzed to ADP, and the P_i_ is released without major rearrangements, causing a further reduction in interactions spanning the ID-OD cleft via the G- and S-loops (loss of five hydrogen bonds between *Pf*ActI and Pγ; not depicted). In a hypothetical model of F-*Pf*ActI, where conformation **2a** is adopted (IV), two interactions formed by the adoption of **1b** (II) are broken, causing a destabilization of the OD in respect to the ID, promoting a filament break. Upon fragmentation and dissociation of the monomer from the newly-formed pointed end, conformation **2a** is retained (V) in the ADP-*Pf*ActI monomer, the nucleotide is exchanged, and conformation **1a** is returned (VI). Changes in the number of interactions on the back face of the monomer (on the inside of the filament) across the ID-OD cleft are highlighted in blue circles. Total interactions (hydrogen bonds, ionic interactions and cation-π interactions) across the ID-OD cleft are 1, 3, 5, 2, 1 in G-Mg-ATP **1a**, G-Mg-ATP **1b**, F-Mg-ADP **1b**, F-Mg-ADP **2a**, and G-Mg-ADP **2a**, respectively, excluding changes caused by loss of Pγ.

## Online Methods

### Mutagenesis

*Pf*ActI mutants were generated by site-directed mutagenesis as described for F54Y and G115A^8^. Mutants A272W, A272C, H74Q, P42Q, E49G, and P42Q/E49G were prepared using similar methods as before, with different primers. All mutants were confirmed by capillary sequencing at the Biocenter Oulu Sequencing Core or at the Center for Medical Genetics and Molecular Medicine, Haukeland University Hospital, Bergen.

### Protein expression and purification

Wild-type and mutant *Plasmodium* actins were purified as described^8,12^. Briefly, insect cell expressed His-tagged *Plasmodium* actins were purified using Ni-NTA affinity chromatography, cleaved with a recombinantly expressed protease (tobacco etch virus protease [TEV] for *Pf*ActI, rhinoviral 3C protease for *Pb*ActII) and passed through a second Ni-NTA column to remove the His-tag and uncleaved protein and finalized by gel filtration over a Superdex 200 column. Mouse gelsolin segment 1 was purified as described^52^ and included in actin samples (where applicable) before gel filtration at a 1.2-fold molar excess.

### Phosphate release assays

P_i_ release was measured using the sensitive 7-diethylamino-3-[N-(2-maleimidoethyl)carbamoyl]coumarin-labeled phosphate-binding protein (MDCC-PBP) biosensor^53,54^ that produces a fluorescence signal upon P_i_ binding. To reduce P_i_, ATP and ADP background, monomeric actins used for P_i_ release assays were pre-treated with DOWEX 1X8 resin (Sigma) equilibrated with G_0_ buffer (10 mM HEPES pH 7.5, 0.2 mM CaCl_2_, 0.5 mM TCEP) for 3 min at 298 K, and further diluted using G_0_ buffer to 1.6-fold higher concentration than that used for measurements. Before initiating the kinetic measurements, components for the different conditions were supplied as 8-fold concentrated stocks such that the desired final concentrations of all components were reached. Final compositions of the three conditions were 10 mM HEPES pH 7.5, 0.2 mM CaCl_2_, 0.5 mM TCEP, 50 μM ATP (Ca condition), Ca condition with added 1 mM MgCl_2_ (Mg condition), and Ca condition with added 4 mM MgCl_2_, 1 mM EGTA, and 50 mM KCl (MgK condition). Fluorescence vs. time data were converted to μM P_i_ by linear interpolation of a standard series and analyzed by linear regression at the linear portion of the kinetic curve. The slope of the regression line was then divided by the protein concentration measured after DOWEX treatment to yield the final rates.

In the presence of Mg and in MgK, α-actin displays an initial lag phase, followed by an exponential P_i_ release curve which, at high concentrations, plateaus close to the upper limit of the linear range of the system (**Supplementary Fig. 1**). We therefore decided to consider only the first two phases for our analyses. We further calculated the activation of P_i_ release by Mg^2+^ and by K^+^ by dividing the Mg rate by the Ca rate in the former and the MgK rate by the Mg rate in the latter. These ratios are a sensitive measure for comparing actins to one another, since they are insensitive to changes in residual nucleotide contamination in the samples. These contaminants are of the order of <10% of the 50 μM ATP added to each reaction. Since the total nucleotide concentration is in a >50-fold excess over the nM-range dissociation constant of ATP to actin^55^, we assume that in the assay, actin is saturated and not affected by small fluctuations in the nucleotide concentration.

### Electron microscopy

*Pf*ActI wild-type and mutant samples were polymerized for 16 h at 298 K at a concentration of 20 μM in F-buffer. Prior to application on carbon-coated 200 mesh Cu-grids (Electron Microscopy Sciences), samples were diluted to a final concentration of 1 μM and immediately applied on the grids. Samples were incubated for 60 s on the grids, dried from the side using pre-wetted Whatman paper, washed with three drops of F-buffer, then stained with 2% uranyl acetate first for 2 s and then for 60 s in a fresh drop, before drying from the side as before and then dried in air. Grids were imaged using a Jeol JEM-1230 microscope operated at 80 kV and with a final pixel size of 1.22 nm.

### Protein crystallization

*Pf*ActI-G1 and *Pb*ActII-G1 complexes in the Mg state were prepared essentially as described^21^, with the exception that CdCl_2_ was replaced by 1 mM MgCl_2_. In some cases, crystals were grown by streak seeding as described^21^, and in others, crystals were obtained directly from optimization screens without seeding. Cryoprotection was achieved by soaking for 5-30 s using the same condition as for the crystallization with a higher precipitant concentration (PEG3350, 22-28%) and PEG400 at 10-20% as the cryoprotectant. Protein buffer components were also included in the cryosolutions at concentrations of 1 mM MgCl_2_, 0.5 mM ADP, and 0.5 mM TCEP for the Mg conditions and 0.2 mM CaCl_2_, 0.5 mM ATP, and 0.5 mM TCEP for the Ca conditions. The pH of the crystallization reservoir buffer (0.1 M Bis-Tris) varied from 5.8 to 6.5. Mg-ADP-*Pf*ActI-G1 crystals were cryoprotected in a solution containing 50 mM potassium phosphate. Mg-ADP-AlF_n_-*Pf*ActI-G1 crystals were prepared by adding a solution of 20% PEG3350, 0.1 M Bis-Tris pH 6.0, 0.2 M KSCN, and 1 mM AlF_n_ solution directly into the drops and incubated for a few minutes before cryoprotection with a solution as described above. The AlF_n_ solution consisted of pre-mixed AlCl_3_ and NaF in a 1:4 molar ratio. The minimum time between data collection from a crystal yielding structures with ATP/ADP mixtures and ADP-only was 6 months for Mg-*Pf*ActI-F54Y crystals, while the time from crystallization to data collection from Mg-*Pb*ActII was only 2 weeks.

### Diffraction data collection, processing, and structure refinement

Crystallization data was collected at 100 K at several beamlines. Mg-ATP/ADP-*Pf*ActI, Mg-ADP-Pr*Pf*ActI, Mg-ADP-F54Y, and Mg-*Pb*ActII were collected at beamline P13 of PETRA III, DESY (Hamburg, Germany), Ca-F54Y and Mg-AlF_n_-F54Y at I24 of Diamond Light Source (Didcot, UK), Mg-F54Y, Mg-H74Q, and Mg-A272W at I04-1 of Diamond Light Source (Didcot, UK), Ca-G115A at ID23-1 of ESRF (Grenoble, France), and Mg-G115A at MX-14.1 of BESSY (Berlin, Germany). Diffraction images were processed using the XDS package^56^. Structure determination and refinement were carried out using programs of the PHENIX suite^57^. Initial phases were found by molecular replacement with PHASER^58^, using the Ca-ATP-*Pf*ActI-G1 structure (PDB ID 4CBU) as the search model for the *Pf*ActI structures and the Ca-ATP-*Pb*ActI-G1 structure (PDB ID 4CBX) for the *Pb*ActII structure. Additionally, MR-SAD using Autosol^59^ was used to reduce model bias in the Mg-ADP-AlF_n_-F54Y structure. Structure refinement was carried out using phenix.refine^60^.

### Principal component analysis

There are two main structural rearrangements recognized in actin: the twistedness of the two main domains (SD1-2 and SD3-4) along an axis that pierces SD1 and SD3 at their respective centers and the openness of the nucleotide binding cleft as a rotation around an axis perpendicular to the twist axis and to the plane of the F-actin monomer. We analyzed 147 unique actin structures found in the protein data bank (PDB) using BIO3D^61^ at resolution ≤ 4Å together with the structures reported here, and found that these movements are captured well by principal component analysis (PCA) in PC1 (twistedness) and PC2 (openness) that contain 78% of total variance (**Supplementary Movie 1**). All actin chains were aligned on a common core before PCA analysis. In a plot of PC1 vs. PC2 (**Fig. 5a-b**), most actin structures cluster at the center of the plot. This large cluster contains all structures reported in this paper. Several outliers to this large cluster form their own distinct groups. Filament structures cluster at low twistedness and average openness, open profilin-actin structures cluster at high openness and average twistedness, free G-actin structures cluster at high twistedness and average openness and finally ADF/cofilin bound actin structures cluster at high twistedness and low openness. We also analyzed *Plasmodium* actin structures as their own set by similar PCA analysis. PC1 and PC2 contain 84% of total variance in this dataset and their trajectories are toned-down versions of the twistedness and openness of the full dataset (**Supplementary Movie 1**). While PC1 in the limited dataset can easily be recognized as the twisting motion of the full dataset (due to the presence of the F-*Pf*ActI model), the opening-closing motion is slightly ambiguous (due to the lack of an open *Pf*ActI model), and is therefore indicated with an asterisk.

### Domain motion analysis

To support the PCA analysis, we measured three parameters of four sets of structures: the domain distance between SD2 and SD4 (d_2-4_), the phosphate clamp distance (b_2_^19^) and the torsion angle defined by all four subdomains (θ). The d_2-4_ distance was measured by a distance between the mass centers of the Cα atoms of residues 35-39, 52-73 for SD2 and 183-269 for SD4. The phosphate clamp distance was measured as the distance between α-carbons of Gly16 and Asp158. The torsion angle θ was measured using the mass centers of α-carbons from all four domains using the residue assignment defined above for SD2 and SD4, as well as residues 6-32, 77-137, 340-366 for SD1 and 140-182, 270-337 for SD3. The models used were (i) Wild-type *Pf*ActI structures in the Ca-ATP, Mg-ATP/ADP, Mg-ADP and F-ADP states, (ii) *Pf*ActI F54Y structures in Ca-ATP, Mg-ADP-AlF_3_, and Mg-ADP, (iii) *Pb*ActII structures in Ca-ATP and Mg-ADP states and (iv) *D. discoideum* actin structures of mutant E205A/R206A/E207A/P109I in Ca-ATP and Mg-ATP and mutant E205A/R206A/E207A/P109A in Ca-ATP and Mg-ADP. Results are presented in **Supplementary Table 4**. For the *D. discoideum* structures, all residue assignments are -1 relative to the numbers presented above. For *Pb*ActII, residue assignments for residue numbers smaller than 232 were -1 relative to *Pf*ActI and others as for *Pf*ActI.

## Acknowledgments

We are grateful for the skillful assistance of Ju Xu and Dr. Juha Vahokoski in constructing some of the *Pf*ActI mutants, Dr. Henni Piirainen for help with the purification of some of them, Dr. Juha Kallio for help with data collection and Arne Raasakka for critical reading of the manuscript. We acknowledge Dr. Martin R. Webb from the Francis Crick Institute for providing the MDCC-PBP plasmid. We thank the Biocenter Oulu Electron Microscopy core facility, in particular Dr. Ilkka Miinalainen, as well as the Molecular Imaging Center, University of Bergen, in particular Dr. Endy Spriet, for assistance with electron microscopy. We also thank the Biocenter Oulu Proteomics and Protein Analysis core facility and Dr. Ulrich Bergmann for assistance with mass spectrometry. We also acknowledge the use of the Diamond Light Source beamlines I24 and I04-1, the European Synchrotron Radiation Facility beamline ID23-1, the European Molecular Biology Laboratory/German Electron Synchrotron beamline P13 on PETRA III, and the Berliner Elektronenspeicherring-Gesellschaft für Synchrotronstrahlung beamline MX-14.1 and thank the facilities for excellent user support during data collection for the structures published here. This work was funded by the Academy of Finland, the Emil Aaltonen Foundation, the Jane and Aatos Erkko Foundation, the Norwegian Research Council, and the Sigrid Jusélius Foundation.

## Author Contributions

E.-P.K. performed all experimental work, with the exception of the following: A.J.L. and L.T. prepared the wild-type and the K270M mutant actins for electron microscopy experiments and performed them and H.H. refined three of the mutant structures (F54Y/Ca-ATP, A272W, and H74Q). E.-P.K. and I.K. designed the study and wrote the manuscript. All authors participated in analyzing the data and read and approved the manuscript.

## Data Availability

The structures have been deposited in the PDB with the codes 6I4D, 6I4E, 6I4F, 6I4G, 6I4H, 6I4I, 6I4J, 6I4K, 6I4L, and 6I4M. All other data that support the findings of this study are available from the corresponding author upon reasonable request.

